# Nociceptor-enriched CAPS1 isoform couples to Ca_V_2.2 channels and underlies inflammatory pain hypersensitivity

**DOI:** 10.64898/2026.05.23.727326

**Authors:** Melissa Daeschner, Emilio Román Mustafa, Andi Morgan, Quiana Tyss Mosley, Eduardo Javier López Soto

## Abstract

How broadly expressed presynaptic proteins acquire the cell-type specificity needed to couple to Ca_V_2 channels in defined neurons, and whether this specialization contributes to behavior, remains poorly understood. We show that alternative splicing of *Cadps*, encoding the vesicle-priming factor CAPS1, generates nociceptor-selective Ca_V_2.2 regulation and underlies inflammatory pain hypersensitivity. Although *Cadps* is comparably expressed across dorsal root ganglion neurons, exon-resolution analysis identified exon 16a, a conserved 23-amino-acid cassette, as selectively enriched in nociceptors. The e16a-containing CAPS1 isoform, but not CAPS1 lacking e16a, enhanced recombinant Ca_V_2.2 currents, whereas disruption of the e16a-encoded sequence blocked this effect and reduced endogenous Ca_V_2.2 currents in nociceptors. Conditional *Cadps* deletion in Trpv1-lineage nociceptors similarly reduced Ca_V_2.2 currents and attenuated capsaicin-evoked inflammatory hypersensitivity while sparing baseline sensitivity, an effect replicated by intrathecal delivery of an exon 16a-derived peptide. Alternative splicing of a presynaptic partner protein thus confers cell-type-specific Ca_V_2.2 coupling and underlies inflammatory pain hypersensitivity.

## Introduction

A central question in synaptic physiology is how broadly expressed presynaptic release machinery acquires cell-type specificity to support distinct physiological functions and behavior. Molecular coupling between voltage-gated Ca_V_2 calcium channels and the presynaptic release machinery—including SNARE proteins, synaptotagmins, RIM, and Munc13—has anchored models of synaptic strength and plasticity for decades (*1, 2*). Yet, the mechanisms by which such coupling becomes cell-type specific, and whether it influences defined behavioral states, remain poorly understood.

Nociceptors, the primary sensory neurons specialized to detect damaging stimuli (*3*), provide an ideal system in which to resolve this question. These neurons rely heavily on Ca_V_2.2 channels as a major source of depolarization-evoked calcium entry to trigger transmitter release in the spinal dorsal horn (*4, 5*). Ca_V_2.2 channels are associated with and regulated by auxiliary subunits (*6, 7*), G-protein-coupled receptors (*8*), and cytosolic proteins including CRMP2 (*9, 10*), which couples to Ca_V_2.2 channels and modulates pain-related behaviors. Moreover, genetic and pharmacological studies further show that Ca_V_2.2 channels become especially important during inflammatory and injury-associated sensitization (*4, 11–14*), a feature that has made Ca_V_2.2 a longstanding therapeutic target for pain control (*15*). Despite the relevance of Ca_V_2.2 channels in nociceptor function, presynaptic release machinery coupled to Ca_V_2.2 channels remain largely uncharacterized at the functional and behavioral level.

Alternative pre-mRNA splicing offers a mechanism by which Ca_V_2.2–release machinery coupling could acquire cell-type specificity. By generating functionally specialized isoforms from broadly expressed genes, alternative splicing expands molecular diversity across the nervous system and allows shared synaptic proteins to be adapted to distinct neuronal contexts (*16, 17*). Ca_V_2.2 channels themselves, encoded by *Cacna1b*, undergo alternative splicing relevant to pain, producing isoforms that influence G-protein signaling and opioid efficacy in nociceptors (*18, 19*). Moreover, alternative splicing is prominent in dorsal root ganglion (DRG) sensory neurons, where it frequently targets broadly expressed neuronal and synaptic genes (*20–22*). These observations raise the possibility that isoform diversity within presynaptic release machinery, not only within Ca_V_2.2 itself, specifies how Ca_V_2.2 channels are functionally coupled in nociceptors.

CAPS1, the calcium-dependent activator protein for secretion 1, is a vesicle priming factor required for calcium-mediated exocytosis in neurons (*23–25*). CAPS proteins were first recognized as essential factors for dense-core vesicle exocytosis and neuropeptide secretion (*24–28*), and later shown to contribute more broadly to synaptic vesicle priming and excitatory neurotransmitter release in mammalian neurons (*23, 29, 30*). CAPS1 facilitates SNARE complex assembly and drives vesicles from a docked to a fusion-ready state (*31–33*)—an obligate step for stimulus-evoked exocytosis (*34*). The gene encoding CAPS1, *Cadps*, is alternatively spliced (*35*), suggesting that CAPS1-dependent priming is molecularly diversified across cellular contexts. Yet the dominant CAPS1 isoform(s) in nociceptors remain unknown; in DRG sensory neurons, CAPS1 is broadly expressed and enriched at synapses, with evidence supporting calcium-dependent synaptic transmission (*36, 37*).

Here, we asked whether alternative splicing of presynaptic release machinery provides a mechanism for specifying Ca_V_2.2 functional coupling in nociceptors and impacting pain behavior. By integrating exon-resolution analyses across independent RNA-seq datasets, we identify Cadps exon 16a, a conserved 23-amino-acid cassette, as selectively enriched in nociceptors despite broad gene-level *Cadps* expression. The e16a-containing CAPS1 isoform, but not the e16a-lacking canonical isoform, enhances recombinant Ca_V_2.2 currents, whereas an e16a-competitive peptide blocks this effect and reduces endogenous Ca_V_2.2 current in nociceptors. Using conditional *Cadps* deletion in Trpv1-lineage nociceptors, we show that endogenous CAPS1 supports ω-conotoxin-GVIA-sensitive Ca_V_2.2 currents and is required for capsaicin-evoked inflammatory hypersensitivity while sparing baseline noxious sensitivity. Critically, intrathecal delivery of the e16a-competitive peptide mimics the behavioral effect of *Cadps* deletion, linking this splice-dependent mechanism to pain behavior. Collectively, our findings establish alternative splicing of presynaptic release machinery as a mechanism that specifies Ca_V_2.2 functional coupling in nociceptors and contributes to inflammatory pain hypersensitivity.

## Results

### Conditional *Cadps* disruption reduces Ca_V_2.2-dependent currents in Trpv1-lineage nociceptors

To examine the role of CAPS1 in nociceptors while avoiding the neonatal lethality associated with constitutive *Cadps* disruption (*38*), we generated Trpv1-lineage-specific conditional *Cadps* knockout mice. Experimental mice carried the *Trpv1^Cre^*allele (*39*) and were homozygous for either the floxed or wild-type *Cadps* allele (Fig. 1A-B). Trpv1-lineage neurons, primarily of C-and Aδ-fibers, include both peptidergic and non-peptidergic nociceptors in mice and account for a major population of DRG neurons involved in the detection of noxious stimuli (*40–45*). In some experiments, the Ai32 Cre-dependent reporter was additionally included (*46*), enabling simultaneous *Cadps* disruption and ChR2-eYFP expression in the targeted Trpv1-lineage population (Fig. 1A-C).

**Figure 1.**
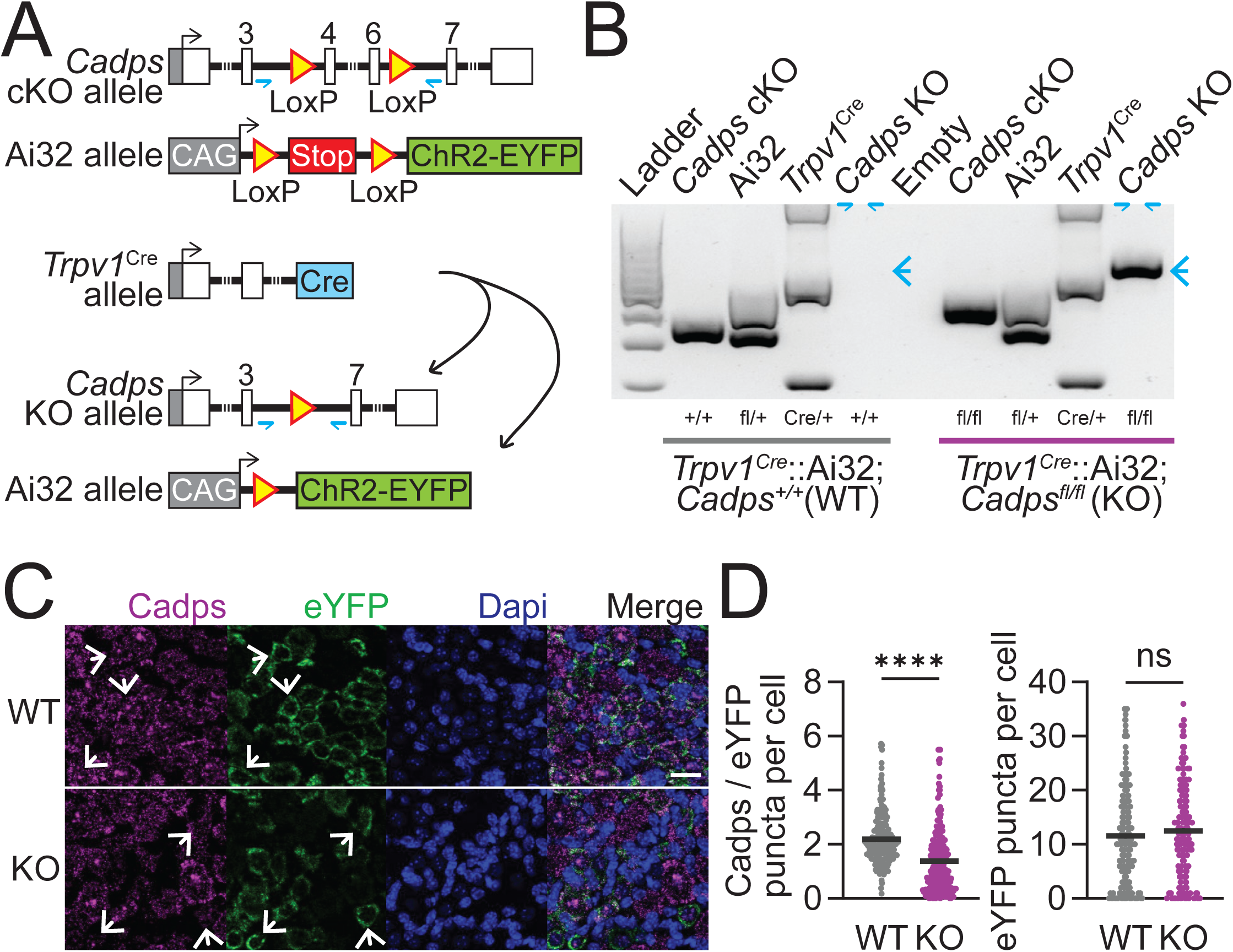
*Cadps* conditional deletion in Trpv1-lineage neurons reduces Cadps expression. Panel A – The Cadps KO allele contains loxP sites flanking the targeted Cadps genomic region, and the Ai32 reporter allele contains a loxP-flanked transcriptional STOP cassette upstream of ChR2-EYFP. In Trpv1Cre-expressing cells, Cre recombines the floxed Cadps allele to generate the Cadps KO allele and excises the STOP cassette in Ai32, thereby activating ChR2-EYFP expression in the same cells. Small half-arrows under the Cadps locus indicate the primer positions used in panel B to detect Cre-mediated recombination. **Panel B –** Representative PCR genotyping of the *Cadps* conditional KO, Ai32, and Trpv1^Cre^ alleles from tail genomic DNA, and of the Cre-recombined Cadps KO allele from DRG genomic DNA, in WT and KO mice. The recombined Cadps KO band, detected with the primers indicated in panel A, is present only in KO mice (arrow), consistent with Cre-dependent recombination of the floxed Cadps allele in Trpv1Cre-expressing cells. **Panel C –** Example confocal images of DRG sections processed for RNA in situ hybridization for Cadps and eYFP transcripts, with DAPI, from WT (top) and KO mice (bottom). eYFP signal is derived from the Ai32 reporter allele marking Trpv1-lineage neurons. Merged images are shown on the right. Arrows indicate example labeled cells. Scale bar, 20 µm. **Panel D –** Quantification of Cadps/eYFP puncta per cell (left) and eYFP puncta per cell (right) in WT and KO DRG sections. Mann-Whitney tests: Cadps/eYFP puncta per cell, WT (n = 251) and KO (n = 291), p < 0.0001 (****); eYFP puncta per cell, WT (n = 313) and KO (n = 348), p = 0.0942 (ns). Outliers were identified using the Tukey interquartile range criterion (Q1 − 1.5×IQR; Q3 + 1.5×IQR) and excluded before statistical analysis. Dots represent individual cells from DRG sections collected from 2 WT and 2 KO mice. Data are shown as mean ± SEM.

PCR genotyping confirmed Cre-dependent recombination of the floxed *Cadps* allele specifically in DRG tissue from postnatal knockout (KO) mice, but not wild-type (WT) controls (Fig. 1B). RNA in situ hybridization showed that Cadps transcript signal was markedly reduced in eYFP-positive Trpv1-lineage neurons from KO mice compared with WT controls, whereas the eYFP-transcript signal across cells was not altered (Fig. 1C-D). These data confirm effective and selective disruption of *Cadps* gene and Cadps expression in the targeted population, with no evidence of loss of the targeted neurons in postnatal mice.

Having validated a viable conditional *Cadps* knockout model, we next asked whether loss of CAPS1 alters voltage-gated calcium channel function in Trpv1-lineage nociceptors. This question was central to our model because Ca_V_2.2 channels provide a major source of depolarization-evoked calcium entry in nociceptors and are strongly implicated in inflammatory pain signaling. We therefore performed whole-cell voltage-clamp recordings from cultured Trpv1-lineage neurons isolated from WT and KO postnatal mice. Nociceptors were identified by eYFP fluorescence and calcium currents were first evoked by depolarizing steps to 0 mV from a holding potential of −80 mV. Under these conditions, peak Ca_V_ current density was significantly reduced in KO neurons relative to WT neurons (Fig. 2A). To resolve the current-voltage (I–V) relationship, we then applied 10 mV incremental depolarizing steps from −60 to +80 mV. Across this voltage range, KO neurons showed a broad reduction in current amplitude compared with WT neurons (Fig. 2B). Analyses of Boltzmann-Ohmic fits to individual I–V curves indicated that this effect is potentially explained by a 33.2 % reduction in maximal conductance (G_max_), whereas the half-activation voltage (V_0.5_), slope factor (*k*), and reversal potential (V_rev_) were not significantly different between genotypes (Fig. 2C). Membrane capacitance was also unchanged (Fig. 2D), supporting the interpretation that recordings were obtained from comparable neuronal populations.

**Figure 2.**
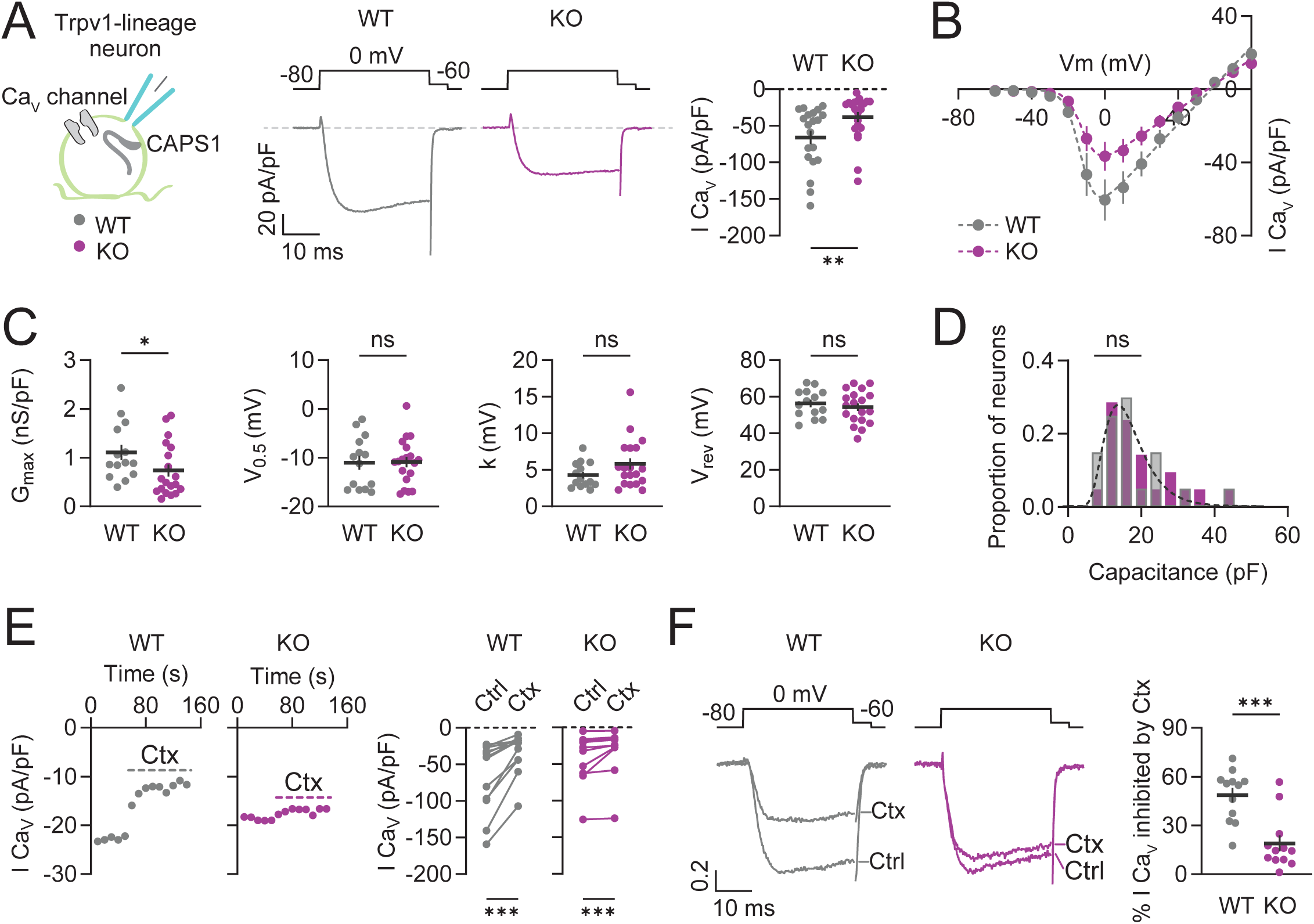
Loss of CAPS1 reduces Ca_V_2.2 current density in Trpv1-lineage nociceptors. **Panel A** – Schematic of the experimental design (left), example traces (middle) and peak current density plot (right) of whole-cell Ca_V_ currents evoked by a depolarization to 0 mV from a holding potential of −80 mV in cultured Trpv1-lineage neurons from WT (n = 20) and KO (n = 22) mice. Mann-Whitney test: p = 0.0050 (**). Dots represent individual cells from five independent DRG cultures. Data are shown as mean ± SEM. **Panel B –** Current-voltage (I–V) relationship curves of whole-cell Ca_V_ currents from cultured Trpv1-lineage neurons from WT (n = 14) and KO (n = 19) mice. Dotted lines represent Boltzmann–Ohmic fits. Data are shown as mean ± SEM. **Panel C –** Summary of biophysical parameters derived from Boltzmann–Ohmic fits to individual I–V curves from panel B for WT (n = 14) and KO (n = 19) neurons: maximal conductance (G_max_), half-activation voltage (V_0.5_), slope factor (k), and reversal potential (V_rev_). Mann-Whitney tests: G_max_: p = 0.0317 (*); Welch’s unpaired t tests: V0.5: p = 0.9328 (ns); Mann-Whitney tests: k: p = 0.2258 (ns); Welch’s unpaired t tests: Vrev: p = 0.4762 (ns). Dots represent individual neurons from five independent DRG preparations. Data are shown as mean ± SEM. **Panel D –** Distribution of whole-cell capacitance values for WT (n = 20) and KO (n = 21) Trpv1-lineage neurons from panel A. Kolmogorov-Smirnov test: p = 0.2480 (ns). One outlier in the KO group was identified using the Tukey interquartile range criterion (Q1 − 1.5×IQR; Q3 + 1.5×IQR) and excluded before statistical analysis. Dotted line indicates the shared lognormal least-squares fit to both datasets; comparison of fits: p = 0.6822 (ns). Data are shown as relative frequency. **Panel E –** Example time courses (left) and peak current density plot (right) of whole-cell Ca_V_ currents evoked at 0 mV before (Ctrl) and during application of ω-conotoxin-GVIA (Ctx, 2 µM) in cultured Trpv1-lineage neurons from WT (n = 12) and KO (n = 12) mice. Wilcoxon matched-pairs signed rank test: WT, p = 0.0005 (***); KO, p = 0.0005 (***). Dots represent individual neurons from five independent DRG cultures. Data are shown as mean ± SEM. **Panel F –** Example current traces (left), normalized to the control peak current, and percentage of whole-cell Ca_V_ current inhibited by ω-conotoxin-GVIA (2 µM) (right) in cultured dissociated Trpv1-lineage neurons from WT (n = 12) and KO (n = 12) mice. Currents were evoked by a step depolarization to 0 mV from a holding potential of −80 mV and recorded before (Ctrl) and during ω-conotoxin-GVIA (Ctx) application. Mann-Whitney test: p = 0.0003 (***). Dots represent individual cells from five independent DRG cultures. Data are shown as mean ± SEM.

We next tested whether the reduction in total Ca_V_ current preferentially reflected loss of Ca_V_2.2 channel activity. Consistent with previous work (*47*), acute application of the selective Ca_V_2.2 blocker ω-conotoxin GVIA (2 μM) substantially reduced Ca_V_ currents in WT Trpv1-lineage neurons (Fig. 2E). ω-Conotoxin GVIA also reduced Ca_V_ currents in KO neurons (Fig. 2E), but its inhibitory effect was significantly smaller in KO neurons than in WT neurons, corresponding to a 62.3 % reduction in the fraction of current inhibited by ω-Conotoxin GVIA (Fig. 2F). These data indicate that CAPS1 is required for normal Ca_V_ channel activity in Trpv1-lineage neurons and suggest that the reduction in total Ca_V_ current reflects a preferential loss of the Ca_V_2.2-dependent activity.

### Cadps exon 16a defines a nociceptor-enriched CAPS1 isoform

Having established that CAPS1 regulates Ca_V_2.2 activity in Trpv1-lineage nociceptors, we next asked whether this function might reflect cell-type-specific features of Cadps expression. RNA in situ hybridization in DRG sections revealed that Cadps transcript signal was broadly distributed across neurons spanning a wide range of soma sizes, with no strong relationship between Cadps transcript density and cell area (Fig. 3A). Because soma area broadly maps onto major sensory neuron classes in the DRG, this pattern suggested that overall Cadps expression is not restricted to a particular subtype. Consistent with this interpretation, reanalysis of published deep RNA-seq datasets from genetically defined sensory neuron populations (*48, 49*) showed that total Cadps expression was broadly comparable across peptidergic and non-peptidergic nociceptors, C-low-threshold mechanoreceptors (C-LTMRs), other LTMR subclasses, and proprioceptors, with only modest subtype-specific differences. Total Cadps expression was also comparable between Trpv1-lineage and non-Trpv1-lineage neurons (Fig. 3B). Thus, overall Cadps abundance does not distinguish nociceptors from other DRG sensory neuron populations.

**Figure 3.**
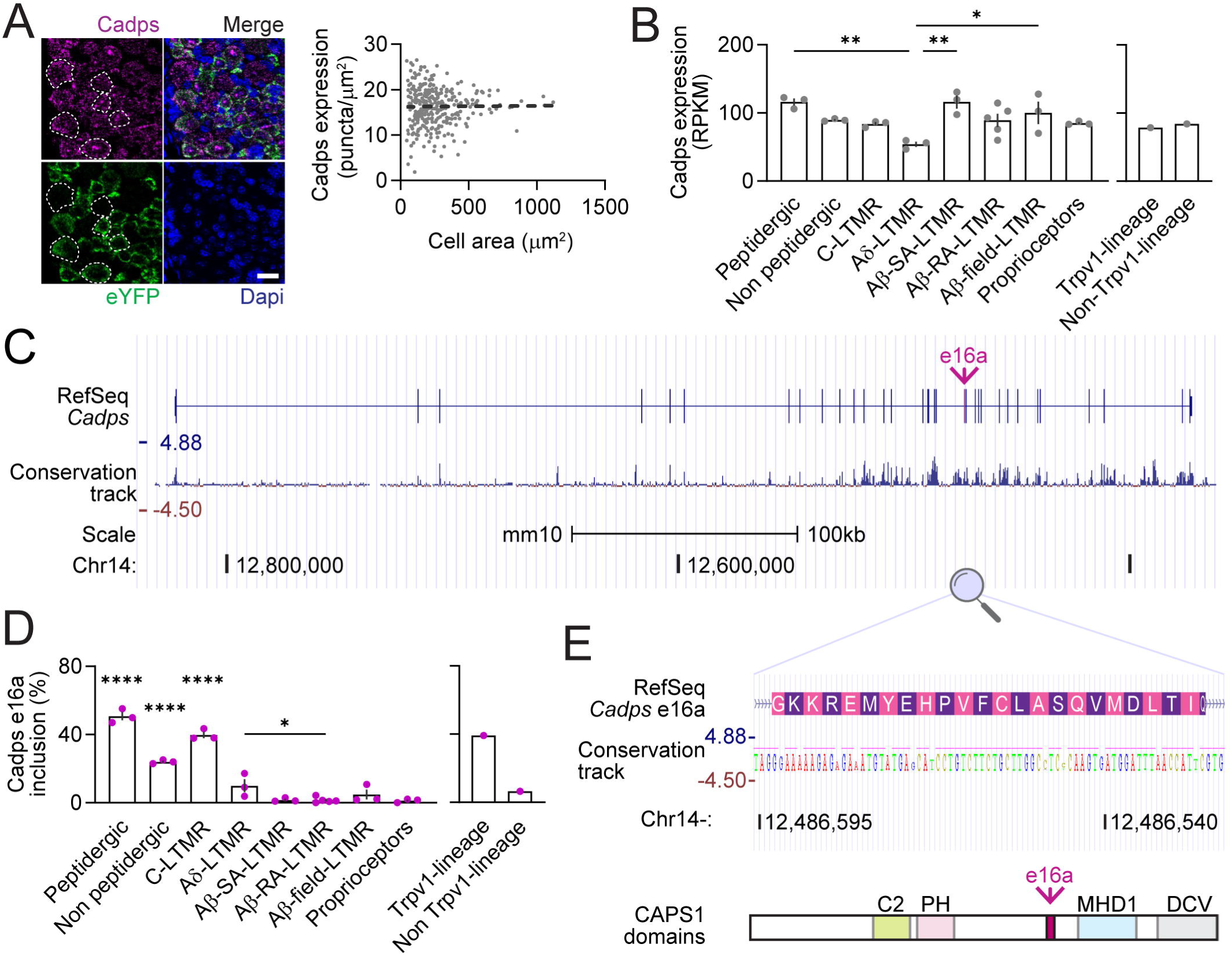
Cadps expression is largely independent of sensory neuron subtype, whereas exon 16a inclusion is enriched in nociceptors and C-LTMRs. **Panel A** – Example confocal images (left) of DRG sections processed for RNA in situ hybridization for Cadps and eYFP transcripts, with DAPI, from WT mice. eYFP signal is derived from the Ai32 reporter allele marking Trpv1-lineage neurons. Merged image is shown at top right. Dashed outlines indicate example ROIs, eYFP positive and negative, used for cell-area measurements. Scale bar, 20 µm. Quantification of Cadps expression per cell versus cell area (right). Dotted line indicates simple linear regression with slope = 0.0002 not significantly different from zero, p = 0.8691. Dots represent individual cells from 2 mice (n = 324). **Panel B –** Cadps expression (RPKM, reads per kilobase per million mapped reads) across genetically defined DRG sensory neuron populations, including peptidergic and nonpeptidergic nociceptors, five low-threshold mechanoreceptor (LTMR) subtypes, and proprioceptors (left) (see Zheng *et al*. 2019 for genetic labeling) and in Trpv1-lineage and non-Trpv1-lineage neurons (right) (see Goswami *et al*. 2014 for genetic labeling) from publicly available RNA-seq datasets (SRP198454; SRP068217). One-way ANOVA: p = 0.0031. Tukey’s multiple comparisons: peptidergic vs. Aδ-LTMR, p = 0.0026 (**); Aδ-LTMR vs. Aβ-SA-LTMR, p = 0.0026 (**); Aδ-LTMR vs. Aβ-field-LTMR, p = 0.0344 (*). Data are shown as mean ± SEM. **Panel C –** UCSC genome browser view of the mouse *Cadps* locus in the mm10 assembly showing the RefSeq gene model, 60-vertebrate basewise conservation track, scale, and chromosome 14 location. Exons are shown as bars, and exon 16a is indicated by the arrow. **Panel D –** Cadps exon 16a inclusion per transcript (%) across genetically defined DRG sensory neuron populations from the same public RNA-seq datasets shown in panel B. One-way ANOVA: p < 0.0001. Holm-Šídák multiple comparisons: peptidergic vs. non-peptidergic, Aδ-LTMR, Aβ-SA-LTMR, Aβ-RA-LTMR, Aβ-field-LTMR, and proprioceptors, p < 0.0001 (****); peptidergic vs. C-LTMR, p = 0.0115 (*); non-peptidergic vs. C-LTMR, p = 0.0003 (***); non-peptidergic vs. Aδ-LTMR, p = 0.0011 (**); non-peptidergic vs. Aβ-SA-LTMR, Aβ-RA-LTMR, Aβ-field-LTMR, and proprioceptors, p < 0.0001 (****); C-LTMR vs. Aδ-LTMR, Aβ-SA-LTMR, Aβ-RA-LTMR, Aβ-field-LTMR, and proprioceptors, p < 0.0001 (****); Aδ-LTMR vs. Aβ-RA-LTMR, p = 0.0340 (*). Data are shown as mean ± SEM. **Panel E –** UCSC genome browser zoom of the Cadps exon 16a region (top) showing the RefSeq sequence and encoded amino acids, 60-vertebrate basewise conservation track, scale, and chromosome 14 location; and CAPS1 domain map (bottom) highlighting the position of exon 16a (arrow) within CAPS1, relative to the previously described C2, pleckstrin homology (PH), Munc13 homology domain 1 (MHD1), and dense-core vesicle–binding (DCV) domains.

We therefore asked whether isoform expression could provide the relevant cell-type specificity. *Cadps* is a large gene with more than 27 exons spanning over 450 Kb in the mouse genome (Fig. 3C), and neuronal genes commonly undergo extensive alternative splicing (*17*). To address this, we repurposed the same deep RNA-seq datasets (*48, 49*) for exon-level analysis using VAST-TOOLS (*17, 50*), a splicing-sensitive pipeline that estimates exon inclusion by aligning and quantifying short sequencing reads to exon-exon junctions. This approach identified six of eight alternative exons within *Cadps* that were expressed across DRG sensory neurons, revealing substantial isoform complexity within the gene. Among these, exon 16a (e16a), located immediately downstream of constitutive exon 16, emerged as a prominent nociceptor-enriched exon. E16a inclusion was substantially higher in peptidergic and non-peptidergic nociceptors and C-LTMRs than in proprioceptors and most low-threshold mechanoreceptor subtypes, and was similarly enriched in Trpv1-lineage relative to non-Trpv1-lineage neurons (Fig. 3D). Genomic analysis further showed that e16a is evolutionarily conserved across vertebrates and encodes an in-frame 23-amino-acid segment inserted into CAPS1 within an intervening region between annotated functional domains (Fig. 3E). These findings suggest that the relevant cell-type specificity in Cadps might not be encoded at the level of overall gene abundance, but rather at the level of alternative splicing, with e16a defining a nociceptor-enriched CAPS1 isoform.

### The nociceptor-enriched CAPS1+e16a isoform increases recombinant Ca_V_2.2 currents

Having identified Cadps e16a as nociceptor-enriched and given that loss of Cadps reduced Ca_V_2.2 currents in Trpv1-lineage nociceptors, we asked whether the e16a-containing CAPS1 isoform (CAPS1+e16a) could directly modulate Ca_V_2.2 channel activity. To test this, we expressed recombinant Ca_V_2.2 channels in tsA201 cells by co-transfecting the pore-forming α1B subunit together with β_3_ and α_2_δ-_1_ auxiliary subunits, either alone or with individual CAPS1 isoforms. Whole-cell voltage-clamp recordings were then used to evoke Ca_V_2.2 currents with depolarizing steps to 0 mV from a holding potential of −100 mV (Fig. 4A). Under these conditions, co-expression of CAPS1+e16a significantly increased peak Ca_V_2.2 current density relative to Ca_V_2.2 expressed alone, whereas co-expression of the e16a-lacking CAPS1 isoform (CAPS1Δe16a) did not produce a detectable increase (Fig. 4A). Analysis of the I–V relationship further showed that the effect of CAPS1+e16a was evident across multiple depolarized voltages (Fig. 4B). Fitting of individual I–V curves indicated that this effect could be primarily explained by an 85.7 % increase in maximal conductance (G_max_), with no significant changes in half-activation voltage (V_0.5_), slope factor (*k*), or reversal potential (V_rev_) across conditions (Fig. 4C). Thus, inclusion of e16a is sufficient to define a functionally distinct CAPS1 isoform that enhances Ca_V_2.2 channel activity.

**Figure 4.**
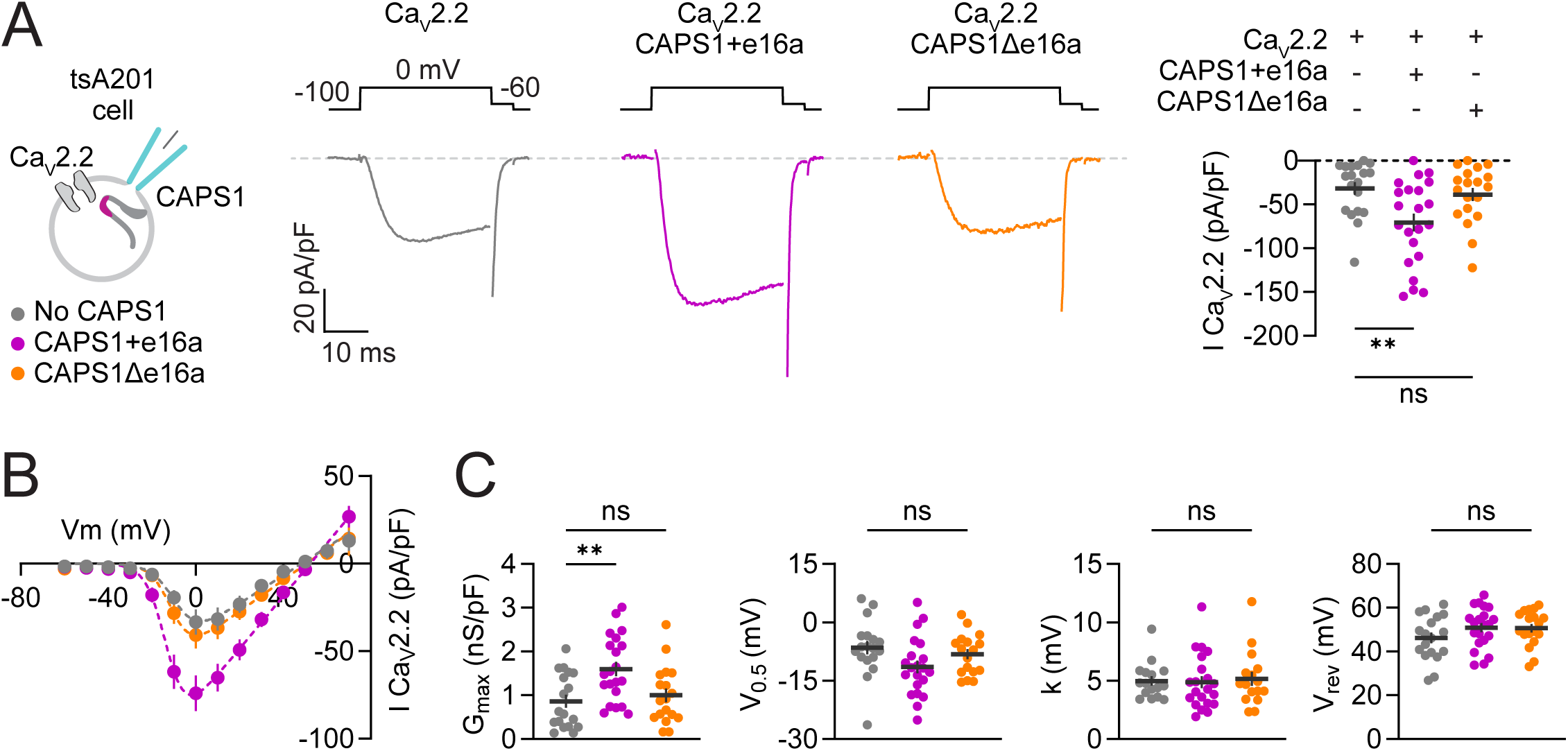
CAPS1+e16a, but not CAPS1Δe16a, selectively enhances Ca_V_2.2 currents in tsA201 cells. Panel A – Schematic of experimental design (left), example traces (middle) and peak current density plot (right) of whole-cell Ca_V_2.2 currents evoked by a depolarization to 0 mV from a holding potential of −100 mV in tsA201 cells expressing recombinant Ca_V_2.2 channels (α1B co-expressed with β3 and α2δ-1 subunits) under three conditions: Ca_V_2.2 alone (n = 19), Ca_V_2.2 + CAPS1+e16a (n = 22), and Ca_V_2.2 + CAPS1Δe16a (n = 19). Kruskal-Wallis test: p = 0.0098. Dunn’s multiple comparisons: Ca_V_2.2 vs. Ca_V_2.2 + CAPS1+e16a, p = 0.0072 (**); Ca_V_2.2 vs. Ca_V_2.2 + CAPS1Δe16a, p = 0.9036 (ns). Dots represent individual cells from four independent transfections. Data are shown as mean ± SEM. **Panel B –** Current-voltage (I–V) relationship curves of whole-cell Ca_V_2.2 currents in tsA201 cells expressing recombinant Ca_V_2.2 channels alone (n = 18), with CAPS1+e16a (n = 21), or with CAPS1Δe16a (n = 18). Dotted lines represent Boltzmann-Ohmic fits. Data are shown as mean ± SEM. **Panel C –** Summary biophysical parameters derived from Boltzmann-Ohmic fits to individual I–V curves shown in panel B for recombinant Ca_V_2.2 channels alone (n = 18), with CAPS1+e16a (n = 21), or with CAPS1Δe16a (n = 18) conditions: maximal conductance (G_max_), half-activation voltage (V_0.5_), slope factor (k), and reversal potential (V_rev_). For G_max_, Kruskal-Wallis test: p = 0.0039. Dunn’s multiple comparisons: Ca_V_2.2 vs. Ca_V_2.2 + CAPS1+e16a, p = 0.0038 (**); Ca_V_2.2 vs. Ca_V_2.2 + CAPS1Δe16a, p > 0.9999 (ns). For V_0.5_, one-way ANOVA: p = 0.0779 (ns). For k, Kruskal-Wallis test: p = 0.7932 (ns). For V_rev_, one-way ANOVA: p = 0.2366 (ns). For G_max_, one outlier in the Ca_V_2.2 alone group and two outliers in the CAPS1Δe16a group were identified using the Tukey interquartile range criterion (Q1 − 1.5×IQR; Q3 + 1.5×IQR) and excluded before statistical analysis. For V_0.5_, one outlier in the CAPS1Δe16a group was identified using the same criterion and excluded before statistical analysis. For k, one outlier in the Ca_V_2.2 alone group and two outliers in the CAPS1Δe16a group were identified using the same criterion and excluded before statistical analysis. Dots represent individual cells from four independent transfections. Data are shown as mean ± SEM.

### The e16a-encoded sequence mediates the CAPS1-dependent increase in Ca_V_2.2 channel activity in nociceptors

If the e16a-encoded sequence contributes to the mechanism by which CAPS1+e16a enhances Ca_V_2.2 currents, then introducing the isolated e16a sequence should competitively disrupt the effect of full-length CAPS1+e16a. To test this, we co-expressed a mRuby3-tagged construct containing the e16a sequence in the recombinant Ca_V_2.2 expression system and recorded channel activity as described above. Expression of the e16a construct alone did not alter baseline Ca_V_2.2 currents (Fig. 5A). In contrast, when co-expressed with full-length CAPS1+e16a, the e16a construct markedly reduced the current increase normally produced by CAPS1+e16a, fully disrupting the functional effect of CAPS1 (Fig. 5A). These results indicate that the e16a-encoded region contributes to the functional coupling mechanism through which CAPS1 enhances Ca_V_2.2 channel activity.

**Figure 5.**
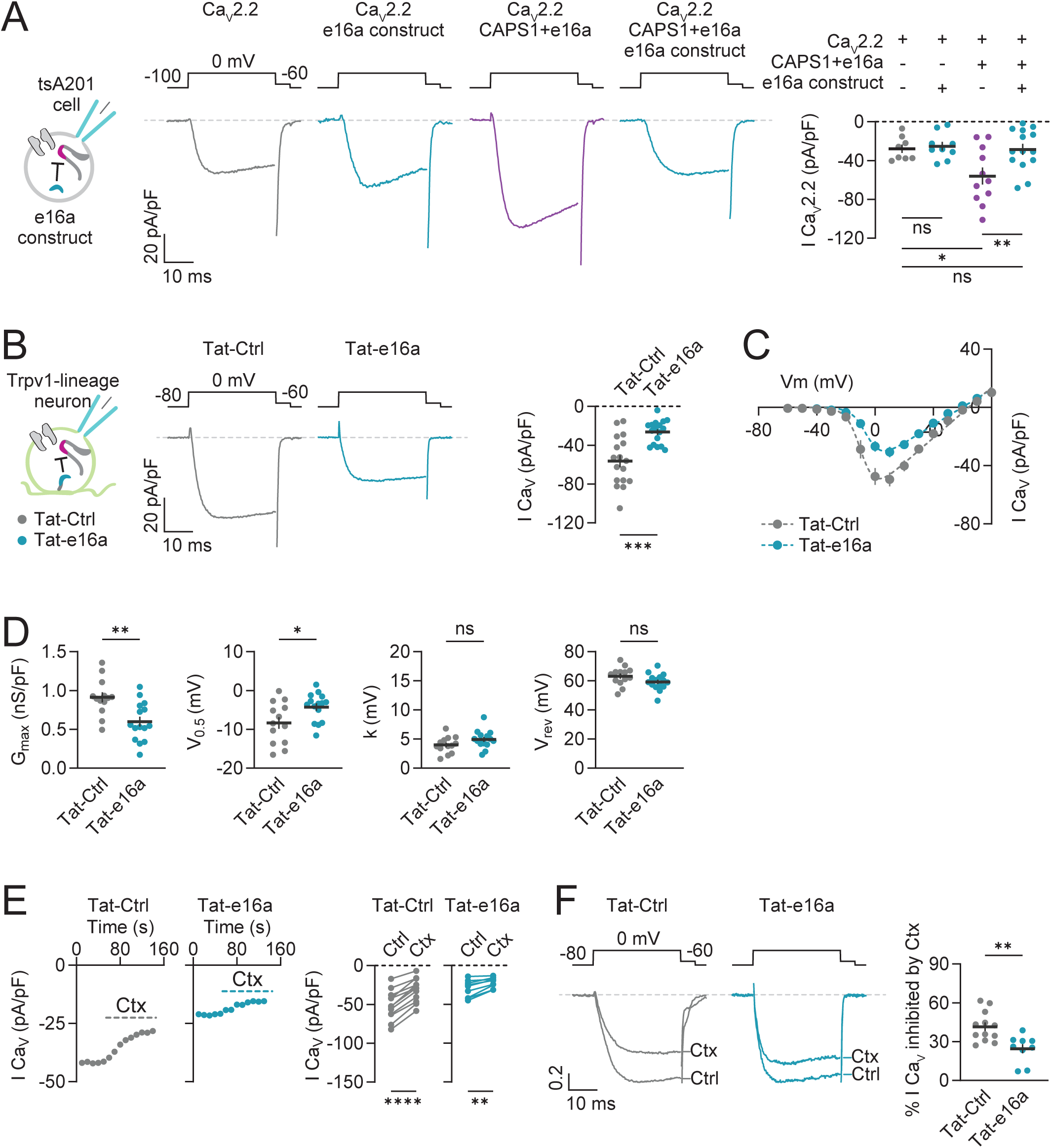
e16a-derived peptide occludes CAPS1-e16a–dependent increase of Ca_V_2.2 and reduces Ca_V_2.2 currents in nociceptors. **Panel A** – Schematic of experimental design (left), example traces (middle) and peak current density plot (right) of whole-cell Ca_V_2.2 currents evoked by a depolarization to 0 mV from a holding potential of −100 mV in tsA201 cells expressing recombinant Ca_V_2.2 channels (α1B co-expressed with β3 and α2δ-1 subunits) under four conditions: Ca_V_2.2 alone (n = 8), Ca_V_2.2 + e16a construct (n = 9), Ca_V_2.2 + CAPS1+e16a (n = 11), and Ca_V_2.2 + CAPS1+e16a + e16a construct (n = 14). One-way ANOVA: p = 0.0049. Sidák multiple comparisons: Ca_V_2.2 vs. Ca_V_2.2 + e16a construct, p = 0.9986 (ns); Ca_V_2.2 vs. Ca_V_2.2 + CAPS1+e16a, p = 0.0254 (*); Ca_V_2.2 vs. Ca_V_2.2 + CAPS1+e16a + e16a construct, p > 0.9999 (ns); Ca_V_2.2 + CAPS1+e16a vs. Ca_V_2.2 + CAPS1+e16a + e16a construct, p = 0.0099 (**). One outlier in the Ca_V_2.2 + e16a peptide group was identified using the Tukey interquartile range criterion (Q1 − 1.5×IQR; Q3 + 1.5×IQR) and excluded before statistical analysis. Dots represent individual cells from four independent transfections. Data are shown as mean ± SEM. **Panel B –** Schematic of experimental design (left), example traces (middle) and peak current density plot (right) of whole-cell Ca_V_ currents evoked by a depolarization to 0 mV from a holding potential of −80 mV in cultured Trpv1-lineage neurons treated for 24 h with 20 µM Tat-Ctrl (n = 17) or Tat-e16a peptide (n = 17). Welch’s unpaired t test: p = 0.0002 (***). Dots represent individual cells from five independent DRG cultures. Data are shown as mean ± SEM. **Panel C –** Current-voltage (I–V) relationship curves of whole-cell Ca_V_ currents from cultured Trpv1-lineage neurons treated for 24 h with 20 µM Tat-Ctrl (n = 13) or Tat-e16a peptide (n = 15). Dotted lines represent Boltzmann-Ohmic fits. Data are shown as mean ± SEM. **Panel D –** Summary biophysical parameters derived from Boltzmann-Ohmic fits to individual I–V curves shown in panel C for Tat-Ctrl (n = 13) and Tat-e16a peptide (n = 15) conditions: maximal conductance (G_max_), half-activation voltage (V_0.5_), slope factor (k), and reversal potential (V_rev_). Welch’s unpaired t tests: G_max_: p = 0.0020 (**); V_0.5_: p = 0.0293 (*); k: p = 0.1072 (ns); V_rev_: p = 0.1048 (ns). One outlier in the Tat-e16a group for k was identified using the Tukey interquartile range criterion (Q1 − 1.5×IQR; Q3 + 1.5×IQR) and excluded before statistical analysis. Dots represent individual cells from five independent DRG cultures. Data are shown as mean ± SEM. **Panel E –** Example time courses (left) and peak current density plot (right) of whole-cell Ca_V_ currents evoked at 0 mV before (Ctrl) and during application of ω-conotoxin-GVIA (Ctx, 2 µM) in cultured Trpv1-lineage neurons treated for 24 h with 20 µM Tat-Ctrl (n = 12) or Tat-e16a peptide (n = 9). Paired two-tailed t tests: Tat-Ctrl: p < 0.0001 (****); Tat-e16a: p = 0.0014 (**). Dots represent individual cells from five independent DRG cultures. Data are shown as mean ± SEM. **Panel F –** Example current traces (left), normalized to the control peak current, and percentage of whole-cell Ca_V_ current inhibited by ω-conotoxin-GVIA (2 µM) (right) in cultured Trpv1-lineage neurons treated for 24 h with 20 µM Tat-Ctrl (n = 12) or Tat-e16a peptide (n = 9). Currents were evoked by a step depolarization to 0 mV from a holding potential of −80 mV and recorded before (Ctrl) and during ω-conotoxin-GVIA (Ctx) application. Welch’s unpaired t test: p = 0.0028 (**). Dots represent individual cells from five independent DRG cultures. Data are shown as mean ± SEM.

We next asked whether this competitive principle extends to native nociceptors. To do this, we designed a Tat-e16a peptide by fusing a 20-amino-acid sequence derived from e16a to the HIV Tat cell-penetrating domain. A scrambled sequence peptide fused to the same Tat domain was used as a control (Tat-Ctrl). Tat-linked peptides have been successfully used as a strategy to competitively disrupt protein-protein interactions, including interactions involving Ca_V_ channel regulatory complexes (*9, 51–53*). Cultured Trpv1-lineage neurons were treated for 24 h with 20 μM Tat-e16a or Tat-Ctrl, after which whole-cell voltage-clamp recordings were performed to measure Ca_V_ currents. Peak Ca_V_ current density was significantly reduced in Tat-e16a-treated neurons compared with Tat-Ctrl-treated neurons (Fig. 5B). The I–V relationship showed reduced current densities across depolarized voltages (Fig. 5C), and analysis of the biophysical fit parameters suggested that the principal effect was a 33.8 % reduction in maximal conductance, accompanied by a 4.1 mV depolarizing shift in half-activation voltage in Tat-e16a-treated cells (Fig. 5D).

ω-conotoxin GVIA application further revealed that Tat-e16a preferentially reduced the Ca_V_2.2-sensitive component. While conotoxin inhibited Ca_V_ currents in both conditions (Fig. 5E), the fraction of current blocked by conotoxin was 40.4 % lower in Tat-e16a-treated neurons relative to Tat-Ctrl-treated (Fig. 5F). Notably, this reduction was similar in magnitude to that observed in KO neurons (Fig. 2F). Thus, competitive disruption of the e16a-encoded sequence is sufficient to decrease endogenous Ca_V_2.2 currents in Trpv1-lineage neurons, supporting the conclusion that CAPS1+e16a is a functionally relevant CAPS1 isoform that promotes enhanced Ca_V_2.2 channel activity in nociceptors.

### *Cadps* disruption produces a selective deficit in nociceptor-driven behavior

The reduction in Ca_V_2.2-dependent currents following *Cadps* disruption raised the possibility that CAPS1 loss may alter nociceptor-dependent behavior. We therefore compared baseline sensory and motor function in Trpv1-lineage *Cadps* KO mice and WT controls and found no detectable differences across these assays. Heat sensitivity measured with the Hargreaves assay was similar between genotypes across a range of radiant heat intensities. In both WT and KO mice, higher stimulus intensities produced shorter paw withdrawal latencies, as expected, indicating preserved behavioral responses to increasing noxious heat (Fig. 6A). Temperature-guided behavior was also unchanged, with both genotypes showing similar avoidance of extreme temperatures in the thermal preference assay (Fig. 6B). Mechanical withdrawal thresholds measured with electronic von Frey testing were not different between groups under baseline conditions (Fig. 6C). Spontaneous locomotor activity was likewise unaffected, as total distance traveled in an activity chamber did not differ between genotypes (Fig. 6D).

**Figure 6.**
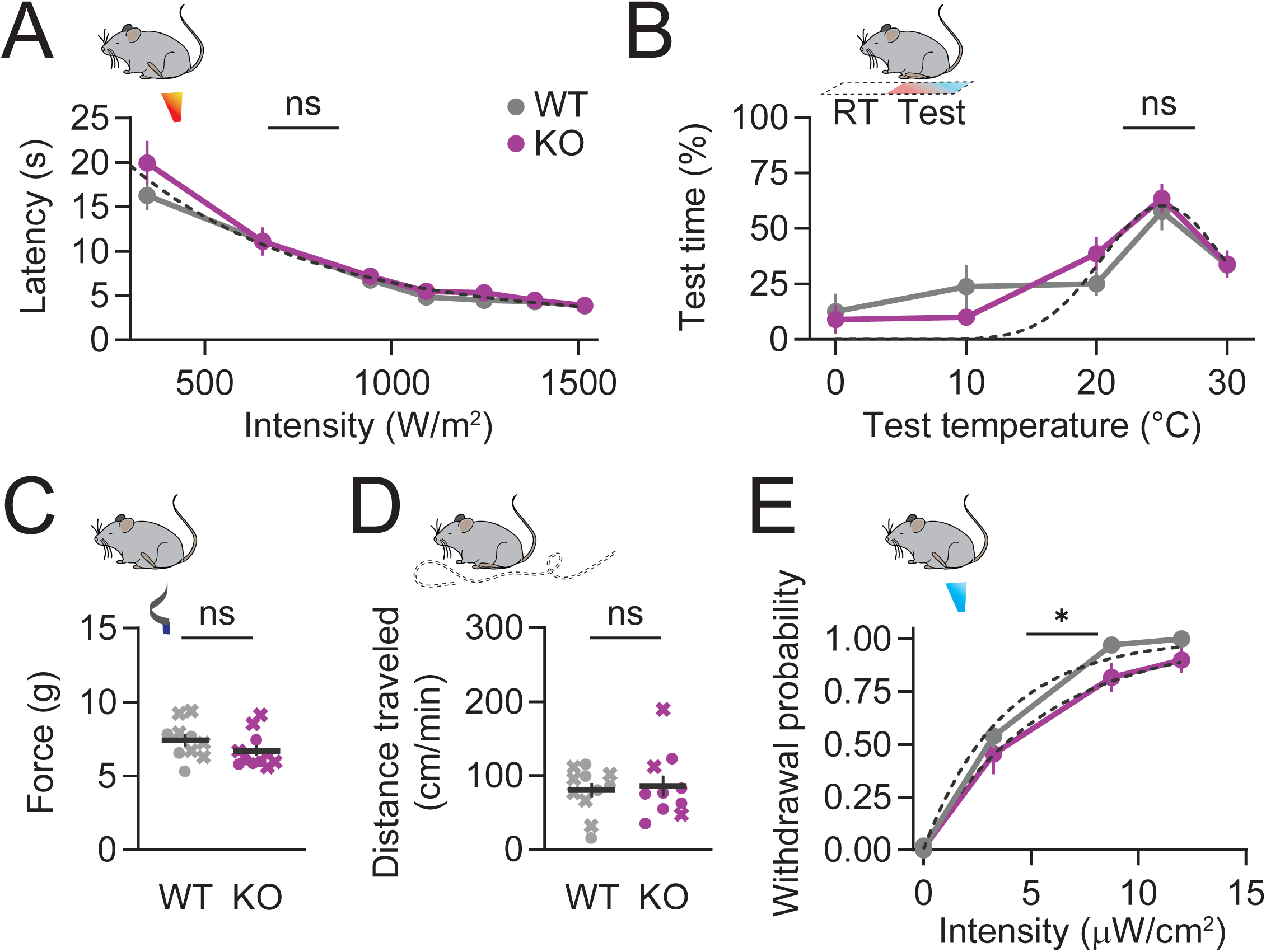
*Cadps* disruption preserves baseline sensory behavior but reduces optogenetically evoked nociceptor responses. **Panel A** – Paw withdrawal latency measured across increasing radiant heat intensities in WT (n = 12) and KO (n = 11) mice. Two-way repeated-measures ANOVA with Geisser-Greenhouse correction: irradiance × genotype, p = 0.4799 (ns); irradiance, p < 0.0001 (****); genotype, p = 0.3794 (ns). Dotted line indicates the shared one-phase decay least-squares fit to both datasets; comparison of fits: p = 0.1578 (ns). Data are shown as mean ± SEM. **Panel B –** Percentage of time spent in the test chamber at each temperature relative to the room-temperature (RT) chamber in WT (n = 13) and KO (n = 15) mice. Two-way repeated-measures ANOVA with Geisser–Greenhouse correction: temperature × genotype, p = 0.3191 (ns); temperature, p < 0.0001 (****); genotype, p = 0.9256 (ns). Dotted lines indicate the shared Gaussian least-squares fit to both datasets; comparison of fits: p = 0.5733 (ns). Data are shown as mean ± SEM. **Panel C –** Mechanical withdrawal threshold in WT (n = 10) and KO (n = 11) mice. Mann–Whitney test: p = 0.1145 (ns). Symbols represent individual mice (crosses indicate males; circles indicate females). Data are shown as mean ± SEM. **Panel D –** Distance traveled in the open field in WT (n = 11) and KO (n = 10) mice. Welch’s unpaired t test: p = 0.7424 (ns). Symbols represent individual mice (crosses indicate males; circles indicate females). Data are shown as mean ± SEM. **Panel E –** Paw withdrawal probability evoked by 3-s blue LED exposure across increasing light intensities in WT (n = 10) and KO (n = 11) mice. Dotted lines indicate the exponential plateau least-squares fits; nonlinear regression comparison of fits: p = 0.0251 (*). Data are shown as mean ± SEM.

To directly evaluate behavior mediated by Trpv1-lineage neurons, we introduced the Ai32 reporter allele into both Cadps KO and WT mice, enabling ChR2-mediated optogenetic stimulation of nociceptors. Mice were exposed to 465 nm light pulses of 3 s duration across ten trials at progressively increasing intensities, and withdrawal probability was recorded as a function of light intensity (Fig. 6E). Both WT and KO mice exhibited intensity-dependent increases in withdrawal probability, confirming effective optogenetic activation of Trpv1-lineage neurons in both genotypes. However, KO mice showed a subtle but consistent reduction in withdrawal probability at higher light intensities compared with WT controls, revealing a deficit in nociceptor-mediated behavioral responses that was not apparent with naturalistic thermal or mechanical stimuli. Because this approach bypasses peripheral sensory transduction and directly drives nociceptor activity, these results indicate that CAPS1 contributes to activation-evoked responses from Trpv1-lineage neurons.

Overall, these findings show that conditional *Cadps* disruption in Trpv1-lineage neurons produces a selective functional deficit rather than a generalized sensory impairment. The absence of baseline changes, together with the emergence of impaired nociceptor-driven behavioral responses under direct activation, supports a specific role for CAPS1 in activity-dependent nociceptor signaling.

### CAPS1 is required for capsaicin-evoked inflammatory hypersensitivity

Because inflammatory sensitization places sustained demand on nociceptor signaling and depends on Ca_V_2.2 channels (*4, 54, 55*), we next tested whether CAPS1 is required during capsaicin-evoked hypersensitivity. Intraplantar capsaicin injection activates TRPV1-expressing peripheral afferents and provides a well-established model of neurogenic inflammation (Fig. 7A). We measured reflexive hind paw withdrawal to heat and mechanical stimulation before and after capsaicin injection, using decreased withdrawal latency and reduced force withdrawal threshold as behavioral readouts of hypersensitivity.

**Figure 7.**
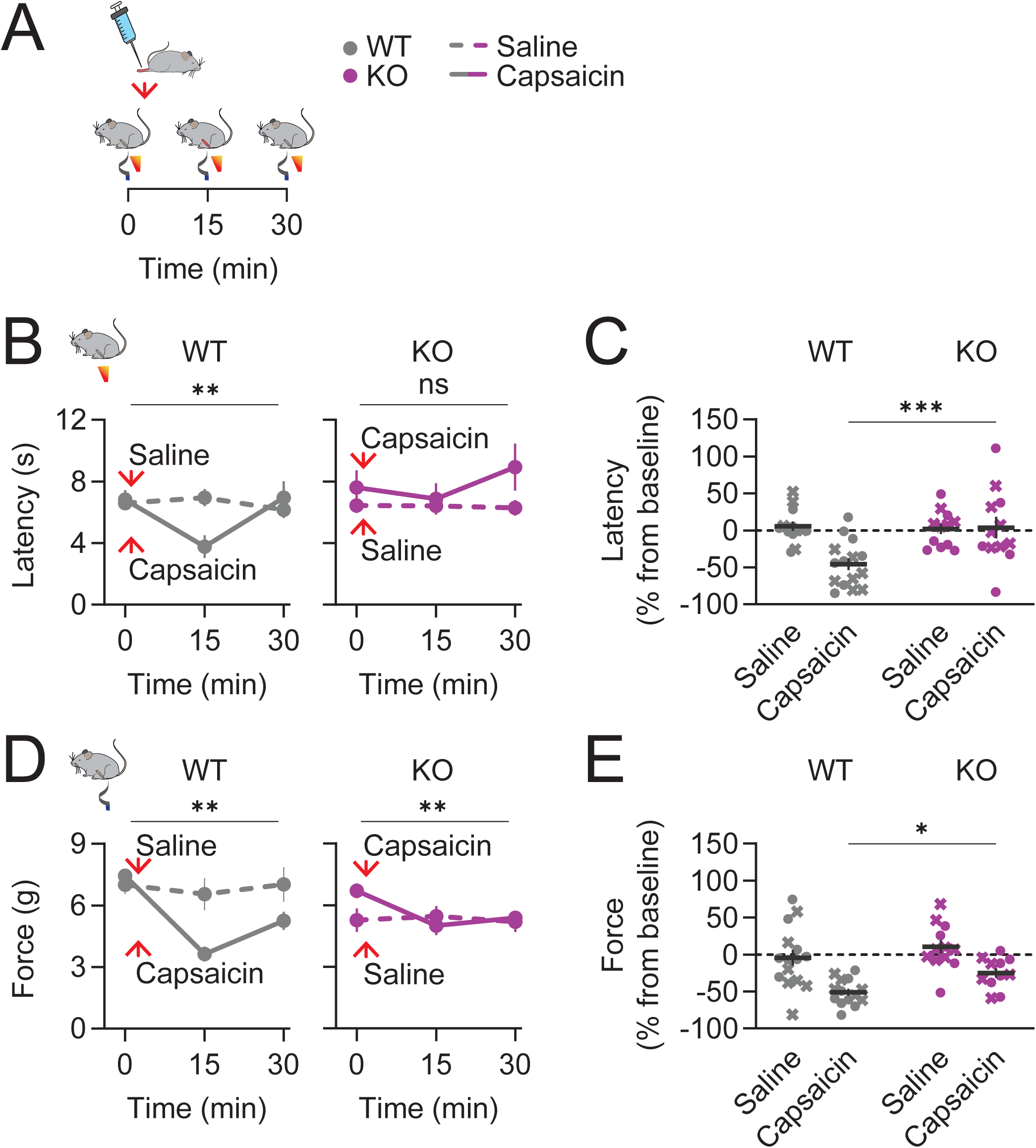
CAPS1 in Trpv1-lineage neurons is required for capsaicin-evoked heat and mechanical hypersensitivity. **Panel A** – Schematic of the experimental design. Saline or capsaicin (20 µL, 0.1 % w/v) was injected intraplantarly in WT or KO mice, and behavioral responses were measured before (0 min) and after injection (15 and 30 min). Red arrow indicates time of injection. **Panel B –** Paw withdrawal latency in response to radiant heat stimuli after intraplantar saline (dotted line) or capsaicin (solid line) in WT (left; saline, n = 13; capsaicin, n = 15) and KO mice (right; saline, n = 12; capsaicin, n = 12). Two-way repeated-measures ANOVAs with Geisser–Greenhouse correction, performed separately within genotype: WT, time × treatment, p = 0.0035 (**); time, p = 0.0482; treatment, p = 0.3659. KO, time × treatment, p = 0.3605 (ns); time, p = 0.4521; treatment, p = 0.1714. Data are shown as mean ± SEM. **Panel C –** Percentage change from baseline in paw withdrawal latency in response to radiant heat stimuli after saline or capsaicin injection in WT and KO mice. Two-way ANOVA: treatment × genotype, p = 0.0064; treatment, p = 0.0111; genotype, p = 0.0183. Šídák multiple comparisons: WT vs. KO, capsaicin, p = 0.0008 (***). Symbols represent individual mice from panel B (crosses indicate males; circles indicate females). Data are shown as mean ± SEM. **Panel D –** Mechanical force required to elicit paw withdrawal after intraplantar saline (dotted line) or capsaicin (solid line) in WT (left; saline, n = 15; capsaicin, n = 14) and KO mice (right; saline, n = 12; capsaicin, n = 11). Two-way repeated-measures ANOVAs with Geisser–Greenhouse correction, performed separately within genotype: WT, time × treatment, p = 0.0016 (**); time, p < 0.0001; treatment, p = 0.0384. KO, time × treatment, p = 0.0077 (**); time, p = 0.0219; treatment, p = 0.4957. Data are shown as mean ± SEM. **Panel E –** Percentage change from baseline in mechanical force required to elicit paw withdrawal after saline or capsaicin injection in WT and KO mice. Two-way ANOVA: treatment × genotype, p = 0.4981; treatment, p < 0.0001; genotype, p = 0.0187. Šídák multiple comparisons: WT vs. KO, capsaicin, p = 0.0357 (*). Symbols represent individual mice from panel D (crosses indicate males; circles indicate females). Data are shown as mean ± SEM.

WT mice treated with 20 µl of 0.1 % (w/v) capsaicin, but not saline, developed robust heat hypersensitivity, reflected by reduced hind paw withdrawal latencies at 15 and 30 min after injection (Fig. 7B). This capsaicin-evoked heat hypersensitivity was strongly attenuated in KO mice, which showed no significant changes in latency across time and treatment (Fig. 7B). Direct comparison of the percent change from baseline in heat withdrawal latency further showed that capsaicin failed to reduce withdrawal latency in KO mice relative to WT mice (Fig. 7C). Capsaicin also produced marked mechanical hypersensitivity, with reduced force thresholds observed after injection in both WT and KO mice, whereas saline did not produce comparable effects in either genotype (Fig. 7D-E). However, the magnitude of this mechanical hypersensitivity was reduced in KO mice compared with WT mice, as the percent change from baseline in mechanical paw withdrawal was significantly attenuated by 51.2 % in KO mice (Fig. 7E). Together, these data show that CAPS1 in Trpv1-lineage neurons is required for the full development of capsaicin-evoked inflammatory hypersensitivity, consistent with a role of CAPS1 in pathways engaged during neurogenic inflammation and sensitized nociceptor activity.

### Intrathecal Tat-e16a peptide attenuates capsaicin-evoked inflammatory hypersensitivity

The genetic and electrophysiological data support a model in which a nociceptor-enriched CAPS1+e16a isoform promotes Ca_V_2.2 channel activity and contributes to inflammatory hypersensitivity. This model predicts that disrupting the e16a-dependent coupling mechanism should attenuate capsaicin-evoked sensitization. To test this, mice received intrathecal injections of 5 µl of 6 µg/kg Tat-e16a or Tat-Ctrl peptide 90 minutes before intraplantar capsaicin treatment, and heat withdrawal latencies were measured at defined time points (Fig. 8A). Intrathecal peptide administration did not itself produce major changes in baseline heat withdrawal latency in either group (Fig. 8B-C). Consistent with this model, Tat-Ctrl-treated mice developed robust capsaicin-evoked heat hypersensitivity, whereas mice receiving Tat-e16a failed to show significant hypersensitivity following capsaicin exposure (Fig. 8B). Direct comparison of the percentage change in withdrawal latency from pre-capsaicin to post-capsaicin confirmed that capsaicin-evoked heat hypersensitivity was significantly attenuated in Tat-e16a-treated mice relative to Tat-Ctrl controls (Fig. 8C).

**Figure 8.**
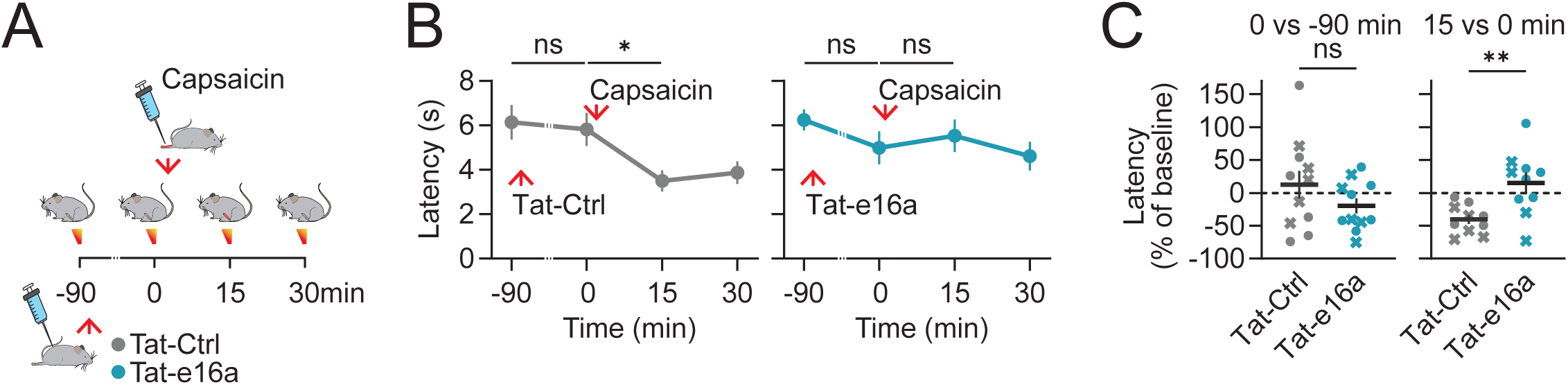
Intrathecal delivery of Tat-e16a peptide attenuates capsaicin-induced heat hypersensitivity. **Panel A** – Schematic of the experimental design. Baseline heat withdrawal latency was measured at −90 min, immediately followed by a 5 µL intrathecal injection of 6 µg/kg Tat-Ctrl or Tat-e16a. Heat withdrawal latencies were reassessed 90 min later (0 min), after which capsaicin (0.1 %, 20 µL) was injected intraplantarly. Withdrawal latencies were subsequently measured at 15 and 30 min post-capsaicin. **Panel B –** Paw withdrawal latency in response to radiant heat stimuli across time, as outlined in panel A, for mice treated with Tat-Ctrl (left, n = 11) or Tat-e16a (right, n = 11). Repeated-measures one-way ANOVAs with Geisser–Greenhouse correction: Tat-Ctrl, p = 0.0185; Tat-e16a, p = 0.2070. Šídák multiple comparisons: Tat-Ctrl, −90 vs 0 min, p = 0.9414 (ns); 15 vs 0 min, p = 0.0123 (*). Tat-e16a, −90 vs 0 min, p = 0.1958 (ns); 15 vs 0 min, p = 0.9677 (ns). Data are shown as mean ± SEM. **Panel C –** Percentage change from baseline in paw withdrawal latency in response to radiant heat stimuli measured post-peptide pre-capsaicin (0 min relative to −90 min; left) and post-peptide post-capsaicin (15 min relative to 0 min; right) in Tat-Ctrl and Tat-e16a treated mice. Welch’s unpaired t tests: 0 vs −90 min, p = 0.3968 (ns); 15 vs 0 min, p = 0.0059 (**). One outlier per group was identified using the Tukey interquartile range criterion (Q1 − 1.5×IQR; Q3 + 1.5×IQR) and excluded before statistical analysis. Symbols represent individual mice from panel B (crosses indicate males; circles indicate females). Data are shown as mean ± SEM.

Finally, we asked whether Tat-e16a treatment resembled the behavioral effect of conditional *Cadps* disruption. The magnitude of the Tat-peptide effect closely mirrored the pattern observed in WT and KO mice after capsaicin. Tat-Ctrl-treated mice showed a percentage change from baseline of −40.28 ± 6.89 % (mean ± SEM; n = 10), matching the −45.87 ± 8.04 % observed in WT mice after capsaicin (n = 15), whereas Tat-e16a-treated mice showed a change of 15.44 ± 15.34 % (n = 10), no different to the 3.79 ± 14.58 % observed in KO mice (n = 12; Welch’s ANOVA, p = 0.0031; Dunnett’s T3 multiple-comparisons: Tat-Ctrl versus WT, p = 0.8396; Tat-e16a versus KO, p = 0.8269). Thus, interfering with the exon 16a-dependent CAPS1 mechanism is sufficient to blunt capsaicin-evoked heat hypersensitivity, mirroring the effect of conditional *Cadps* disruption.

## Discussion

Over the past 30 years, studies of Ca_V_2.2 channels and synaptic protein interactions have established a mechanistic framework for how the calcium source and calcium-sensing mechanisms are linked to evoke transmitter release (*1, 2, 56*). These interactions explain important aspects of synaptic plasticity but often lack defined neuronal-type specificity or behavioral correlates. By considering cell-specific alternative splicing within the release machinery, we demonstrate that the CAPS1 isoform containing the alternatively spliced e16a enhances Ca_V_2.2 channel activity in Trpv1-lineage nociceptors, and that this isoform-specific mechanism contributes to capsaicin-evoked inflammatory hypersensitivity.

Many large synaptic genes undergo extensive alternative splicing, generating isoform diversity that is increasingly recognized as a major source of functional specialization across neuronal types (*57, 58*). However, the physiological meaning of most splice isoforms remains unresolved. *Cadps* e16a provides a concrete example of how a short, evolutionarily conserved alternatively spliced exon can confer functional specificity to a presynaptic effector protein in a defined sensory neuron population. The finding that small-diameter sensory neuron populations, including peptidergic and non-peptidergic nociceptors and C-LTMRs, show high e16a inclusion, whereas most mechanoreceptor and proprioceptor populations showed low or minimal inclusion, suggests that e16a may serve as a molecular determinant that adapts CAPS1 function to the release demands of specialized sensory neurons. This pattern aligns with a growing appreciation that cell-type-specific neuronal function is often encoded in alternatively spliced exons (*18, 59, 60*) and parallels prior demonstrations that sensory neurons rely on splicing programs—including those governed by RBFOX1, CTCF, and other neural factors—to tune sensory function and pain-related plasticity (*20, 22, 61*).

Our findings extend the functional repertoire of CAPS1 upstream vesicle priming (*23, 25, 62*) and position it as a regulator of Ca_V_2.2 channel activity in nociceptors. Ca_V_2 channels are known to interact with components of the release machinery, including SNARE proteins (*63, 64*) and active-zone scaffolds such as RIM and Munc13 (*65, 66*), and the synprint region of Ca_V_2 channels provides a docking site for SNARE binding that can influence both channel gating and release efficacy (*67, 68*). Our data suggest that nociceptor-enriched inclusion of e16a in CAPS1 adds another layer to this organization by favoring molecular configurations that enhance Ca_V_2.2 function. The principal biophysical effect of CAPS1+e16a—an increase in apparent maximal conductance estimated from Boltzmann–Ohmic fits to peak I–V relationships—is consistent with a mechanism that increases the number of functional channels available at the membrane, stabilizes channel complexes, or enhances channel open probability, rather than one that primarily alters voltage-dependent gating. Whether the e16a-encoded segment contacts the Ca_V_2.2 complex directly or acts through an intermediate scaffold remains to be defined.

Inclusion of e16a in CAPS1 in nociceptors may stabilize a molecular configuration that reinforces presynaptic Ca_V_2.2 function and allows transmitter release to scale with sensory demand. Because transmitter release depends supralinearly on local calcium entry (*56, 69, 70*), even moderate e16a-dependent changes in Ca_V_2.2 channel activity at nociceptor terminals could produce large effects on release probability, particularly for the peptidergic pathways engaged during inflammation. The behavioral phenotype is consistent with this model. Conditional *Cadps* disruption preserved baseline thermal preference, mechanical thresholds, and graded heat responses, but attenuated capsaicin-evoked hypersensitivity and reduced optogenetically driven nociceptor responses at high stimulation intensities. This profile—limited effects at baseline with stronger deficits under sensitized or high-activity conditions—parallels observations from Ca_V_2.2-null and Ca_V_2.2-targeted pharmacological studies (*4, 12, 54*) and reinforces the idea that CAPS1 and Ca_V_2.2 contribute to a shared nociceptor signaling pathway. It is also consistent with the established role for Ca_V_2.2-driven neuropeptide release, including CGRP and substance P, in driving neurogenic inflammation and central sensitization (*3, 71, 72*). Although we did not directly measure neurotransmitter or neuropeptide release, the convergence between reduced Ca_V_2.2-sensitive current and attenuated inflammatory hypersensitivity suggests that CAPS1+e16a may be particularly important for release pathways recruited during capsaicin-evoked sensitization.

The peptide competition experiments provide a functional test of the splice-isoform-specific mechanism in native nociceptors and *in vivo*. Peptide-based approaches have been used successfully to disrupt Ca_V_ channel-associated signaling complexes (*9, 51, 73*), supporting the use of Tat-e16a as a tool to interfere with an exon-defined regulatory site. In cultured Trpv1-lineage nociceptors, Tat-e16a reduced the conotoxin-sensitive component of Ca_V_ current, and intrathecal delivery attenuated capsaicin-evoked heat hypersensitivity to a degree that closely matched the *Cadps* KO phenotype. Importantly, this intervention did not alter baseline heat withdrawal, suggesting preferential engagement of sensitized signaling. These findings do not define the precise binding interface between the e16a-encoded sequence and the Ca_V_2.2 channel complex, nor do they distinguish whether Tat-e16a acts through direct competitive displacement, allosteric disruption, altered trafficking, or actions at central terminals. Nevertheless, the convergence of heterologous sufficiency and the ability of Tat-e16a to reproduce key electrophysiological and behavioral features of *Cadps* disruption supports the conclusion that e16a is functionally important for CAPS1-dependent regulation of Ca_V_2.2 channels. More broadly, these results suggest that isoform-specific regulatory interfaces within nociceptor presynaptic machinery may offer a more selective route to modulating Ca_V_2.2-dependent inflammatory pain than broad channel blockade, which is limited by the widespread functions of Ca_V_2.2 channels in non-nociceptor populations (*15*).

In summary, alternative splicing of the *Cadps* gene generates a nociceptor-enriched CAPS1 isoform, CAPS1+e16a, that enhances Ca_V_2.2 channel activity and contributes to capsaicin-evoked inflammatory hypersensitivity. The e16a-dependent mechanism can be competitively disrupted to reduce Ca_V_2.2 currents in native nociceptors and attenuate inflammatory pain behavior. This mechanism is not apparent from gene-level expression and emerges only through exon-resolution analysis, illustrating how isoform-specific coupling between a vesicle-priming factor and presynaptic calcium channels can specialize broadly expressed release machinery for the demands of a defined sensory population and link this specialization to a behavioral output. Beyond its specific implications for Ca_V_2.2 regulation and nociceptor signaling, this work supports a broader framework in which alternative splicing of presynaptic effector proteins represents a tunable layer of synaptic regulation, one that can be experimentally manipulated and, in principle, has therapeutic potential.

## Materials and methods

### Animals

All animal procedures were approved by the Institutional Animal Care and Use Committee at North Carolina State University and conformed to NIH guidelines. Mice were group-housed on a 12 h light/12 h dark cycle with *ad libitum* access to food and water. Both male and female mice were used, and experimental groups were matched for sex and age when possible. Mice were ≥7 weeks old for behavioral studies and postnatal days 21–28 for *ex vivo* recordings. All lines were maintained on a C57BL/6 background. *Trpv1*^Cre^ mice (B6.129-*Trpv1*^tm1(cre)Bbm^/J; JAX 017769) and Ai32 reporter mice (B6.Cg-*Gt(ROSA)26Sor*^tm32(CAG-COP4*H134R/EYFP)Hze^/J; JAX 024109) were obtained from The Jackson Laboratory and the *Cadps*^flox^ mice from GemPharmatech (C57BL/6JGpt-*Cadps*^em1Cflox^/Gpt; T038365). Conditional knockout (KO) mice were obtained by crossing *Cadps^fl/fl^*males with *Cadps^fl/fl^; Trpv1^Cre/+^* females. Cre-negative littermates served as wild-type (WT) controls. For experiments requiring optogenetic stimulation or fluorescent identification of *Trpv1*-lineage neurons, the Ai32 reporter was introduced by crossing to produce *Cadps^fl/fl^; Trpv1^Cre/+^; Ai32/+* (KO) or *Cadps^+/+^; Trpv1^Cre/+^; Ai32/+* (WT) mice.

### Genotyping

Tail biopsies were collected at postnatal day 7 and stored at −20 °C until processing. Genomic DNA was extracted and genotyped using the Azura Mouse Genotyping Kit (AZ-1851) according to the manufacturer’s protocol. *Trpv1^Cre^*, Ai32, and *Cadps^fl^* alleles were genotyped using the primers listed in The Jackson Laboratory protocols 35042 and 28710 and the GemPharmatech protocol for mouse line T038365, respectively. PCR products were resolved by agarose gel electrophoresis.

### PCR confirmation of Cre-dependent *Cadps* recombination in DRG

Lumbar DRGs were dissected, and genomic DNA was isolated using the Monarch® Genomic DNA Purification Kits (76339-770) according to the manufacturer’s protocol. Cre-dependent recombination of the floxed *Cadps* allele was assessed by PCR using primers flanking the excised cassette: forward, 5′-ATC GCA TTG TCT GAG TAC GTG-3′; reverse, 5′-CAG GAT TCT GTC TGT GAT GGT TTC G-3′. PCR reactions were performed using VWR Fast HiFi DNA Polymerase 2X Master Mix (76620-584) with the following cycling conditions: initial denaturation at 98 °C for 2 min; 35 cycles of 98 °C for 20 s, 63 °C for 30 s, and 72 °C for 2 min; followed by a final extension at 72 °C for 5 min. PCR products were resolved by agarose gel electrophoresis, and recombination-specific products were confirmed by Sanger sequencing.

### DRG RNA in situ hybridization (RNAscope), imaging and quantification

Postnatal day 21–28 mice were deeply anesthetized with isoflurane for approximately 5 min and euthanized by cervical dislocation followed by decapitation. The dorsal skin was opened to expose the vertebral column, which was then removed, isolated, and bisected longitudinally. The bisected spinal column was transferred to ice-cold phosphate-buffered saline (PBS; Gibco, 20012-027) in a dish maintained on ice. Lumbar DRGs were identified and dissected under cold PBS, then fixed in 4 % paraformaldehyde (PFA; Electron Microscopy Sciences, 15710) for 24 h. Fixed DRGs were dehydrated through a graded ethanol series, paraffin-embedded, and sectioned at 5 µm by the Histology Research Core Facility (Department of Cell Biology and Physiology, University of North Carolina at Chapel Hill). Tissue processing was performed using a Sakura Tissue-Tek VIP 5 Tissue Processor.

RNA *in situ* hybridization was performed with the RNAscope Multiplex Fluorescent V2 Detection Kit (ACD Bio, 323110) following the manufacturer’s instructions. Briefly, sections were deparaffinized, rehydrated, subjected to antigen retrieval (15 min, 100 °C), digested with protease (30 min, 40 °C), and hybridized with target probes for 2 h at 40 °C. Probes used were Mm-*Cadps* (ACD Bio 882451) and EYFP-C2 (ACD Bio, 312131-C2). Signal was amplified and developed with TSA Plus Fluorescein (Akoya NEL741001KT) and TSA Plus Cyanine 3 (Akoya, NEL744001KT) at 1:1500. Slides were mounted in Fluorogel II with DAPI (Electron Microscopy Sciences, 17985-50).

Images were acquired on a Zeiss LSM 880 confocal microscope with Airyscan (Cellular and Molecular Imaging Facility, North Carolina State University) using the 20X and 40X (water immerse) objectives Image analysis was performed manually in Fiji (ImageJ, v1.54p) using regions of interest (ROIs) to define the cell area based on signal from the eYFP and *Cadps* channels and quantified with the Analyze Particles function to determine the number of puncta per ROI and the corresponding soma cross-sectional area. For each cell, *Cadps*/eYFP puncta ratios and per-cell eYFP puncta counts were computed and compared between WT and KO genotypes.

### Cell-type–resolved RNA-seq and alternative splicing analysis

Cell-type–resolved expression and alternative splicing of *Cadps* were analyzed from publicly available deep RNA-seq datasets of genetically defined mouse DRG sensory neuron populations: Zheng et al. (GEO/SRA accession SRP198454) (*48*), spanning peptidergic and non-peptidergic nociceptors, C-, Aδ-, Aβ-SA-, Aβ-RA-, and Aβ-field low-threshold mechanoreceptors (LTMRs), and proprioceptors; and Goswami et al. (SRA accession SRP068217) (*49*), comprising Trpv1-lineage and non-Trpv1-lineage neurons.

Raw paired-end FASTQ files were processed using VAST-TOOLS v2.5.1 (*17*) with the mouse VASTDB release vastdb.Mm2.23.06.20. Paired-end reads were aligned using vast-tools align with the species option -sp mm10, expression quantification enabled with --expr, the species-specific VASTDB directory supplied with --dbDir, and four processing threads (-c 4). VAST-TOOLS aligns reads to predefined exon–exon and exon–intron junction libraries from the species-specific VASTDB annotation to quantify annotated alternative splicing events. Sample-level outputs were combined using vast-tools combine to generate exon-inclusion and gene-expression tables.

Gene-level *Cadps* expression was extracted as cRPKM values, corrected reads per kilobase per million mapped reads, from the VAST-TOOLS expression output. Exon-level inclusion was quantified as percent spliced-in (PSI) from the VAST-TOOLS INCLUSION_LEVELS_FULL output table. Events with insufficient read support were excluded according to the VAST-TOOLS quality annotations. For exon-skipping events, only PSI values with VLOW or higher coverage scores were retained, corresponding to the default VAST-TOOLS minimum coverage class for quantified events; events below this threshold were treated as not quantified. Six of the eight annotated alternative exons within *Cadps* were detectable in DRG datasets, and percentage e16a inclusion was reported per population.

The mouse Cadps genomic locus was visualized using the UCSC Genome Browser (*74*) on the GRCm38/mm10 assembly. Gene structure was based on NCBI RefSeq annotations, and evolutionary conservation was assessed using the 60-vertebrate basewise conservation track calculated by phyloP (*75*).

### Primary DRG culture

DRGs were dissected as described above and collected from all spinal levels under a dissecting microscope into ice-cold Hank’s balanced salt solution (HBSS; Gibco 24020117) containing 0.5 mg/mL collagenase type I (Gibco, 17100017) and 0.0025 % (w/v) trypsin type II-S (Sigma, T7409). Tissue was digested at 37 °C for 45 min with gentle inversion every 15 min, then triturated 30 times with a fire-polished Pasteur pipette. The cell suspension was layered onto 12 % BSA (Cell Signaling, 9998S) in DMEM (Gibco, 10569-010) and centrifuged at 1000 rpm for 15 min. The pellet was resuspended in Neurobasal-A medium supplemented with B-27, GlutaMAX, and penicillin/streptomycin, recentrifuged at 1000 rpm for 3 min, filtered through a 100-µm strainer (Corning, 431752), and counted on a hemocytometer. Cells were plated at 200–500 cells per coverslip on poly-L-lysine-coated (Fisher, A005C) glass coverslips in 24-well plates and cultured at 37 °C, 5 % CO_2_. Recordings were performed 48–72 h after plating. For peptide-treatment experiments, neurons were cultured for 24 h, then incubated with 20 µM Tat-e16a or Tat-Ctrl peptide for an additional 24 h prior to recording. *Trpv1*-lineage neurons were identified by ChR2-eYFP fluorescence (Ai32 reporter).

### Plasmids and molecular cloning

Plasmids encoding Ca_V_2.2 α_1_B (Addgene, 62574), Ca_V_α_2_δ-_1_ (Addgene, 26575), and Ca_V_β_3_ (Addgene, 26574) were gifts from Diane Lipscombe. The pcDNA3-EGFP plasmid (Addgene, 13031) was a gift from Doug Golenbock. CAPS1 cDNA constructs encoding CAPS1+e16a or CAPS1Δe16a were custom-generated by GENEWIZ using the mouse Cadps expression plasmid OriGene MC224325 as the parental construct. These constructs were engineered to either include or lack alternative exon 16a and were sequence-verified before use.

A minimal mRuby3-tagged e16a construct encoding the e16a sequence was generated for competition experiments. Briefly, the *Cadps* e16a was PCR-amplified using primers LSLab55 5′-TAAGCAGAATTCCTGAGTATGCCAAAATTGAAG-3′ and LSLab57 5′-CTAGAGGATCCTCATTGAATGGTTAAATCCATC-3′, which introduced EcoRI and BamHI restriction sites, respectively. The reverse primer also introduced an immediate stop codon at the C terminus of the amplified sequence. The PCR product and mRuby3-C1 vector (Addgene, 127808; a gift from Salvatore Chiantia) were digested with EcoRI-HF (NEB R3101S) and BamHI-HF (NEB R3136S). Digested products were gel-purified using a Qiagen gel extraction kit (Qiagen 28704) and ligated using the NEB Quick Ligation Kit (NEB M2200S). The ligation product was transformed into One Shot TOP10 chemically competent E. coli (Thermo Fisher Scientific, C404003), plasmid DNA was isolated using the QIAprep Spin Miniprep Kit (Qiagen, 27104), and the final construct was sequence-verified by Plasmidsaurus.

### tsA201 cell culture and transfection

tsA201 cells, kindly provided by Dr. Lipscombe (Brown University), were maintained in DMEM (Gibco, 10569-010) supplemented with 10 % fetal bovine serum (Gibco, A5670401) at 37 °C and 5 % CO₂. Cells were passaged at ∼80 % confluence using 0.0 5% trypsin-EDTA (Gibco, 25200-056).

For transfection, cells were seeded in 24-well plates and transfected at ∼70 % confluence using Lipofectamine 2000 (Invitrogen, 11668019) in Opti-MEM reduced-serum medium (Gibco, 31985070), according to the manufacturer’s instructions. For Ca_V_2.2 current recordings, cells were transfected with Ca_V_2.2 α_1_B, α_2_δ-_1_–HA, Ca_V_β_3_, and the indicated CAPS1 and or e16a construct at a 1:1:1:1:1 molar ratio. Empty pcDNA3.1 vector was used to balance the total amount of DNA across control conditions. Each well received 0.6 µg total DNA, including 0.05 µg pcDNA3-EGFP to identify transfected cells. Cells were used 24 h after transfection.

### Whole-cell patch-clamp electrophysiology

Whole-cell voltage-clamp recordings were performed at room temperature using an Axopatch 200B amplifier (Molecular Devices), digitized at 10 kHz, and low-pass filtered at 10 kHz with pCLAMP 11 software (Molecular Devices). Patch borosilicate glass pipettes (Warner Instruments G85150T-3) were pulled to a resistance of 4–6 MΩ. Series resistance was monitored throughout each recording; cells with R_s_ > 6 MΩ were excluded, and remaining series resistance was compensated to 80 % with a 10 µs lag. Leak currents were subtracted online using a P/−4 protocol. For recombinant Ca_V_2.2 recordings in tsA201 cells, the external solution contained (in mM): 2 CaCl_2_, 4 MgCl_2_, 10 HEPES, and 135 choline chloride (pH 7.4, CsOH). For native Ca_V_ recordings in cultured *Trpv1*-lineage neurons, the external solution contained (in mM): 135 TEA-Cl, 2 CaCl_2_, 4 MgCl_2_, and 10 HEPES (pH 7.4, TEA-OH). The pipette (internal) solution contained (in mM): 134 CsCl, 10 EGTA, 1 EDTA, 10 HEPES, and 4 Mg-ATP (pH 7.2, CsOH). tsA201 cells were held at −100 mV to maximize Ca_V_2.2 channel availability before depolarization, whereas native DRG neuron recordings were performed from a holding potential of −80 mV. Peak Ca_V_ currents were evoked by 25 ms step depolarizations to 0 mV applied every 10 s. Current–voltage (I–V) relationships were generated by 10 mV depolarizing steps from −60 to +80 mV. I–V curves from individual cells were fit with a Boltzmann–Ohmic equation, as previously described for Ca_V_2.2 currents (*76*): y = g(x - E)/(1 + e^(-(x - V)/k)) where *y* is the current amplitude, *x* is the test potential, *g* is maximal conductance (G_max_), E is the reversal potential (V_rev_), V is the half-activation voltage (V_0.5_), and k is the slope factor. Whole-cell capacitance (C_m_) was used as an estimate of cell surface area. For pharmacological isolation of Ca_V_2.2-mediated currents, ω-conotoxin GVIA (2 µM; Sigma 343781) was applied to the bath while currents were continuously monitored at 0 mV from −80 mV holding potential. The percentage of Ca_V_ current inhibited by ω-conotoxin GVIA was calculated as [1 − (I_Ctx / I_Ctrl)] × 100, where I_Ctrl is the peak current before toxin application and I_Ctx is the peak current after ω-conotoxin GVIA application.

### Tat-e16a peptide design and synthesis

*Cadps* e16a is located at chr14:12,486,525–12,486,593 (GRCm38/mm10), is 69 nucleotides long, and encodes a 23-amino-acid sequence, GKKREMYEHPVFCLASQVMDLTI. To generate a cell-permeable peptide competitor, we selected a 20-amino-acid sequence from e16a, GKKREMYEHPVFCLASQVMD, excluding the final three residues to improve aqueous solubility while preserving most of the exon-specific coding sequence. This e16a-derived sequence was fused at the N-terminus to the HIV-1 Tat protein transduction domain, <u>YGRKKRRQRRR</u>, to enhance membrane permeability, resulting in Tat-e16a: <u>YGRKKRRQRRR</u>GKKREMYEHPVFCLASQVMD, with a predicted molecular weight of 3.91 kDa. A control peptide, Tat-Ctrl, was generated by scrambling the e16a-derived portion while preserving the Tat sequence and peptide length: <u>YGRKKRRQRRR</u>MYHLADKSCKQVGPRFEVME, also with a predicted molecular weight of 3.91 kDa. Peptide specificity was assessed using NCBI BLASTP. The N-terminal Tat cell-penetrating sequence was excluded from the query, and only the e16a-derived or scrambled cargo peptide sequence was searched. BLASTP searches were performed against the mammalian RefSeq protein database with parameters adjusted for short peptide sequences. Searches used an Expect threshold of 0.05, word size 5, and the BLOSUM62 matrix. Significant matches were defined a priori as alignments with E-value < 1 × 10⁻³ and ≥80 % sequence identity over ≥50 % of the query sequence. Under these criteria, the e16a-derived peptide matched mammalian CAPS1/Cadps sequences, including mouse CAPS1, with 100 % query coverage and 100 % sequence identity, and no significant non-CAPS1 mammalian protein matches were detected. The scrambled control peptide showed no significant mammalian protein matches under the same criteria. Peptides were synthesized by GenScript with HPLC purity ≥95.0 % and reconstituted in sterile UltraPure water for *in vitro* and *in vivo* experiments.

### Behavioral assays

#### General

Behavioral testing was performed during the light phase on mice aged ≥7 weeks. Animals were habituated to the testing apparatus for at least 20 min prior to each assay. Experimenters were blinded to genotype and treatment during data collection and analysis; each assay was performed by at least two experimenters. Sex- and age-matched WT and KO littermates were randomized. A positive response was scored only when paw withdrawal was accompanied by paw flicking, licking, guarding, or grimace-scale features.

#### Heat sensitivity

Radiant heat sensitivity was measured with the Hargreaves apparatus (Ugo Basile, 37570). Mice were placed in individual Plexiglas enclosures on a glass platform and allowed to habituate for at least 20 minutes. An infrared (IR) heat source was directed onto the plantar surface of the hind paw, and the latency to paw withdrawal was recorded. Three trials per hind paw were averaged with ≥1 min between trials. A 30 s cut-off was used to avoid tissue injury. For baseline heat sensitivity, withdrawal latencies were measured across seven IR intensities corresponding to a calibrated irradiance range of 344.94–1518 W/m^2^, as measured with a IR power meter (Hojila, LH-131). For capsaicin-evoked hypersensitivity experiments, an IR intensity corresponding to 1383.9 W/m^2^ was used.

#### Thermal preference

Thermal preference was assessed using a Thermal Place Preference/Two-Temperature Choice Nociception Test apparatus (Bioseb BIO-T2CT). Mice were allowed to freely explore two adjoining plates, with one plate maintained at a neutral temperature of 30 °C and the other set sequentially to a test temperature of 0, 10, 20, 25, 30 °C. Each test lasted 3 min, and the position of the neutral plate was alternated across trials to control for side bias. Mouse position was tracked using the manufacturer’s T2CT software, and thermal preference was calculated as the percentage of time spent on the test plate relative to the neutral plate.

#### Mechanical sensitivity

Mechanical withdrawal thresholds were measured using a Dynamic Plantar Aesthesiometer/electronic von Frey apparatus (Ugo Basile, 37550). Mice were placed on an elevated mesh platform and allowed to habituate for at least 20 min. A blunt-tipped filament was applied to the plantar surface of the hind paw at a ramp rate of 1 g/s, with a 20 g cut-off. The force eliciting paw withdrawal was recorded. Three trials per hind paw were averaged, with ≥1 min between trials.

#### Optogenetic activation of Trpv1-lineage neurons

WT and KO mice carrying an Ai32 allele (*Cadps^fl/fl^; Trpv1^Cre/+^; Ai32/+* (KO) or *Cadps^+/+^; Trpv1^Cre/+^; Ai32/+* (WT) mice) were placed on an elevated glass platform habituated before testing. A 465 nm LED light source (Thorlabs LEDD1B, M470F4) was directed onto the plantar hind paw at three intensities (3.254, 8.756, and 12.016 µW/cm^2^; calibrated with Hojila LH-139 blue light meter). Each intensity was delivered ten times per mouse in random order using 3 s light pulses, with ≥1 min between trials. Withdrawal probability at each intensity was calculated as the number of trials in which the mouse responded divided by the total number of trials.

#### Locomotor activity

Spontaneous locomotor activity was assessed using the Thermal Place Preference/Two-Temperature Choice Nociception Test apparatus (Bioseb BIO-T2CT). Mice were allowed to freely explore the two adjoining plates for 3 min, with both plates maintained at 30 °C. Mouse position was tracked using the manufacturer’s T2CT software, and total distance traveled during the session was recorded.

#### Capsaicin-evoked neurogenic inflammation

Mice were briefly anesthetized with isoflurane and received an intraplantar injection of 20 µL of 0.1 % (w/v) capsaicin (Sigma PHR1450) into the plantar surface of one hind paw. Capsaicin was first dissolved in 100 % ethanol and then diluted in sterile saline to the final concentration. Control mice received an equal volume of sterile saline as vehicle. Heat and mechanical sensitivity were measured at baseline and 15 and 30 min after injection on the injected paw as described above. Triplicate measurements per paw were averaged at each time point. Percent change from baseline was calculated as ((T_post − T_baseline) / T_baseline) × 100.

#### Intrathecal peptide delivery

Mice were deeply anesthetized with isoflurane delivered through a SomnoSuite Small Animal Anesthesia System (Kent Scientific). Hair over the lumbar spine was clipped and the L4–L5 interspace was identified by palpation. A 33-gauge needle on a 100 µL Nanofil syringe (World Precision Instruments NF100, IO-KIT) coupled to a UMP3T-1 microinjection pump was inserted intrathecally; correct placement was confirmed by a brisk tail flick. Five µL of 6 µg/kg Tat-e16a or Tat-Ctrl peptide per mouse in sterile saline were injected at 2500 nL/min, and the needle was held in place for 60 s after the end of infusion before slow withdrawal. Mice recovered in clean cages. Heat withdrawal latencies were measured immediately before peptide injection (baseline, −90 min), 90 min after peptide injection (post-peptide, pre-capsaicin; 0 min), and 15 and 30 min after subsequent intraplantar capsaicin injection. Experimenters were blinded to peptide identity throughout.

## Statistical analysis

All data are reported as mean ± SEM, with individual data points shown and sample sizes indicated in figure legends. In scatter plots, males are shown as crosses and females as circles. Sample sizes were chosen based on pilot data and prior work using comparable preparations; no formal power calculation was performed. Experiments were not powered to detect sex-specific effects; therefore, sex was not included as an independent biological variable in the statistical models. Data from both sexes were pooled after inspection showed no obvious sex-specific divergence in the direction of the effects. Statistical analyses were carried out in GraphPad Prism (v11). Distributions were assessed for normality by the Shapiro–Wilk test, and the appropriate parametric (unpaired or paired two-tailed Student’s *t* test with Welch’s correction; one- or two-way ANOVA with Geisser–Greenhouse correction for repeated measures, followed by Šídák’s, Tukey’s, Dunnett’s, or Holm–Šídák multiple-comparisons tests) or non-parametric (Mann–Whitney *U*, Wilcoxon matched-pairs signed rank, Kruskal–Wallis with Dunn’s multiple comparisons, Kolmogorov–Smirnov) test was selected for each comparison; the specific test applied is indicated in each figure legend. Welch’s ANOVA with Dunnett’s T3 multiple-comparisons was used for the cross-cohort comparison of capsaicin-evoked percent change in withdrawal latency across the four experimental groups (WT, KO, Tat-Ctrl, Tat-e16a; Fig. 8). Non-linear fits (Boltzmann–Ohmic, one-phase decay, Gaussian, exponential plateau, lognormal) were compared between groups using the extra-sum-of-squares *F* test. Outliers were identified using the Tukey interquartile-range criterion (values < Q1 − 1.5 × IQR or > Q3 + 1.5 × IQR) and excluded prior to analysis where indicated in the corresponding figure legends. Significance was set at p < 0.05; exact p values are reported in figure legends.

## Acknowledgments

The authors thank members of the Lopez Soto laboratory and colleagues at North Carolina State University for scientific input and helpful discussions throughout this study. The authors also thank Dr. Jesica Raingo for critical reading of the manuscript and Dr. Allison Dickey from the NCSU Bioinformatics Research Center for technical assistance with the bioinformatic analysis. The authors thank Dr. Diane Lipscombe, Dr. Doug Golenbock, and Dr. Salvatore Chiantia for providing plasmids used in this study. This work was supported by the National Institutes of Health (NIH; R00NS116123 to E.J.L.S.) and a College of Veterinary Medicine - NC State University intramural award to E.J.L.S. The authors acknowledge the use of the Cellular and Molecular Imaging Facility (CMIF) at North Carolina State University, which is supported by the State of North Carolina and the National Science Foundation.

## Author contributions

E.J.L.S. and M.D. designed the experiments and led the project. M.D., Q.T.M. and E.J.L.S. performed and analyzed behavioral experiments. M.D., E.R.M., and E.J.L.S. performed and analyzed patch-clamp recordings Q.T.M and E.J.L.S. performed the splicing analyses. A.M. contributed to cloning, mouse-line validation, and technical assistance. E.J.L.S. conceived the study and wrote the manuscript. All authors critically reviewed and approved the final manuscript.

## Data availability

All source data underlying the figures are provided with this manuscript. Public RNA-seq datasets reanalyzed in this study are available from the NCBI Sequence Read Archive under accession numbers SRP198454 (*48*) and SRP068217 (*49*). Plasmids generated in this study will be deposited at Addgene.

